# AVMoments-EEG: A Large-Scale EEG Dataset of Naturalistic Audiovisual Event Perception

**DOI:** 10.64898/2026.09.17.751663

**Authors:** Shuning Tang, Zitong Lu, Leiting Li, Dongwei Li

**Author notes:** Please address correspondence to and. These authors contributed equally to this work (Shuning Tang, Zitong Lu).

## Abstract

Understanding how the human brain processes dynamic audiovisual events requires datasets that combine naturalistic stimulation, high temporal resolution, and systematic experimental manipulation of sensory information. Here, we introduce AVMoments-EEG, the first large-scale, high-temporal-resolution human EEG dataset designed to characterize neural responses to naturalistic audiovisual events. Across 10 participants, we recorded 64-channel scalp EEG during free viewing of 3-second naturalistic video clips derived from the Audiovisual Moments in Time dataset. The stimulus set comprised 896 training videos (16 event types × 56 videos per type), each presented three times, and 64 held-out test videos (16 types × 4 videos per type), each presented 40 times under three conditions: (1) intact audiovisual input, (2) intact visual input with scrambled audio, and (3) intact auditory input with scrambled visual input. This design yielded 10,368 trials per participant and combines broad stimulus coverage with highly repeated test stimuli. AVMoments-EEG enables temporally resolved investigation of naturalistic event processing and modality-specific audiovisual contributions, while providing a benchmark for video-to-EEG encoding, EEG-based decoding, and brain–model alignment. The dataset offers a resource for studying dynamic event perception, multisensory processing, and the temporal correspondence between human neural responses and artificial neural network representations.

## Background & Summary

In everyday life, humans continuously understand the surrounding world by integrating what they see and hear. Unlike static object recognition, natural event perception requires the brain to extract information from continuously changing sensory inputs and integrate these signals into coherent event representations^1-3^. Capturing these rapidly evolving processes therefore requires neural measurements with high temporal resolution, for which electroencephalography (EEG) is particularly well suited. Large-scale EEG datasets with diverse naturalistic stimuli further enable systematic characterization of these dynamics and provide benchmarks for neural encoding, decoding, and brain–model alignment. In recent years, representative datasets, including the ImageNet-based EEG dataset constructed by Spampinato et al.^4^, THINGS-EEG^5^, and THINGS-EEG2^6^, have substantially advanced research on visual representations, neural encoding, and brain-signal decoding. However, these datasets primarily rely on static, unimodal visual stimuli and therefore provide limited access to the temporal and multisensory computations involved in natural event perception. Therefore, it is crucial to construct a large-scale, multimodal EEG dataset under naturalistic dynamic event stimuli.

Such a dataset could support research at multiple levels. First, dynamic videos allow researchers to examine how the brain extracts action information from temporally continuous sensory input, identifies event boundaries, and organizes local changes into stable event representations. Second, dynamic audiovisual stimuli can help researchers reveal how sensory information from different modalities is extracted, how visual and auditory signals influence one another, and how it is integrated into unified perceptual and semantic representations. Third, a large number of stimuli and densely repeated measurements can support video-to-EEG encoding models and EEG-to-video decoding models, helping researchers rigorously evaluate the correspondence between computational features and neural dynamics. Finally, when combined with richly annotated naturalistic videos, such datasets can also be used to compare the temporal correspondence between artificial neural networks and the human brain during the processing of action, event, and semantic information.

In recent years, studies have attempted to record brain activity using dynamic naturalistic stimuli, such as movies and natural videos. Existing video EEG datasets have recorded brain activity while participants watched movies or video clips, but many were designed for specific applications such as affective processing or memory. For example, FACED^7^ recorded EEG while participants watched emotion-eliciting videos, providing an important resource for research on affective processing and neural responses under naturalistic stimulation. The Essex EEG Movie Memory Dataset^8^ recorded EEG while participants watched movie clips and was used to study memory for and recognition of naturalistic videos. Compared with large-scale image-based EEG resources, these datasets generally contain fewer distinct video exemplars and do not provide the combination of category-balanced event stimuli, dense stimulus repetitions, and controlled audiovisual manipulations needed for systematic event-level modeling. In contrast, short-video fMRI datasets, such as the BOLD Moments Dataset^9^ and the Human Action Dataset^10^, have expanded the number and category range of dynamic event stimuli. However, the limited temporal resolution of fMRI makes it difficult to track the rapidly unfolding neural dynamics involved in natural event processing. Therefore, there remains a need for a neural dataset that combines naturalistic dynamic audiovisual events, dense repeated measurements, explicit modality manipulations, and millisecond-level temporal resolution. Together, these limitations leave a gap for an EEG resource that combines large-scale naturalistic event sampling, dense repetitions, controlled audiovisual manipulations, and millisecond temporal resolution.

To address this gap, we developed AVMoments-EEG. The Audiovisual Moments in Time (AVMIT) dataset^11^ provides an appropriate stimulus foundation for such a resource. Derived from Moments in Time dataset^12^, AVMIT contains carefully curated 3-s naturalistic audiovisual even clips with clear event labels and strong correspondence between visual and auditory information. We used its controlled stimulus set of 960 videos spanning 16 event categories to construct our AVMoments-EEG. We recorded 64-channel scalp EEG from 10 participants during free viewing of these naturalistic audiovisual events across eight experimental sessions, yielding 10,368 trials per participant. The dataset combines broad stimuli coverage with densely repeated test measurements: 896 distinct training videos were each presented three times, whereas 64 held-out test videos were each presented 40 times under three conditions—intact audiovisual input, intact visual input with scrambled audio, and intact auditory input with scrambled visual input. All data are organized according to the Brain Imaging Data Structure (BIDS)^13^, and preprocessing code and rearranged stimulus-level ERP data are provided to facilitate reuse. By combining naturalistic dynamic events, controlled audiovisual manipulations, a large training stimulus set, and highly repeated test stimuli, AVMoments-EEG provides a resource for investigating the temporal dynamics of event perception and audiovisual processing, developing and benchmarking neural encoding and decoding models, and examining temporally resolved alignment between artificial neural networks and the human brain.

## Methods

### Participants

Thirteen healthy adults were recruited to participate in the experiment and received monetary compensation. One participant (“sub-009”) did not complete all experimental sessions and was therefore excluded from the final dataset. Two additional participants (“sub-006” and “sub-012”) were excluded following the trial-level quality-control procedure applied during construction of the stimulus-level ERP data, because one or more video stimuli had no valid trials remaining after artifact rejection, preventing the construction of a complete stimulus-level ERP dataset. The final dataset therefore comprised 10 participants (6 females and 4 males), with a mean age of 22.4 years (SD = 2.17, range = 20–26 years).

All included participants reported normal or corrected-to-normal vision, normal hearing, and no history of neurological or psychiatric disorders. The study was approved by the ethics committee of Beijing Normal University, and written informed consent was obtained from all participants before the experiment.

### Stimuli

The intact audiovisual stimuli were derived from the Audiovisual Moments in Time (AVMIT) dataset, which was developed from the original Moments in Time (MIT) video dataset (Fig. 1B). MIT contains a large collection of 3-s naturalistic video clips depicting a broad range of actions and dynamic events. AVMIT further annotated and curated a subset of these videos to identify clips in which the labeled event was clearly represented in both the visual and auditory streams, thereby providing a stimulus set with stronger audiovisual correspondence and greater experimental control. Detailed information about the construction and annotation of AVMIT is available in the original publication.

**Fig. 1.**
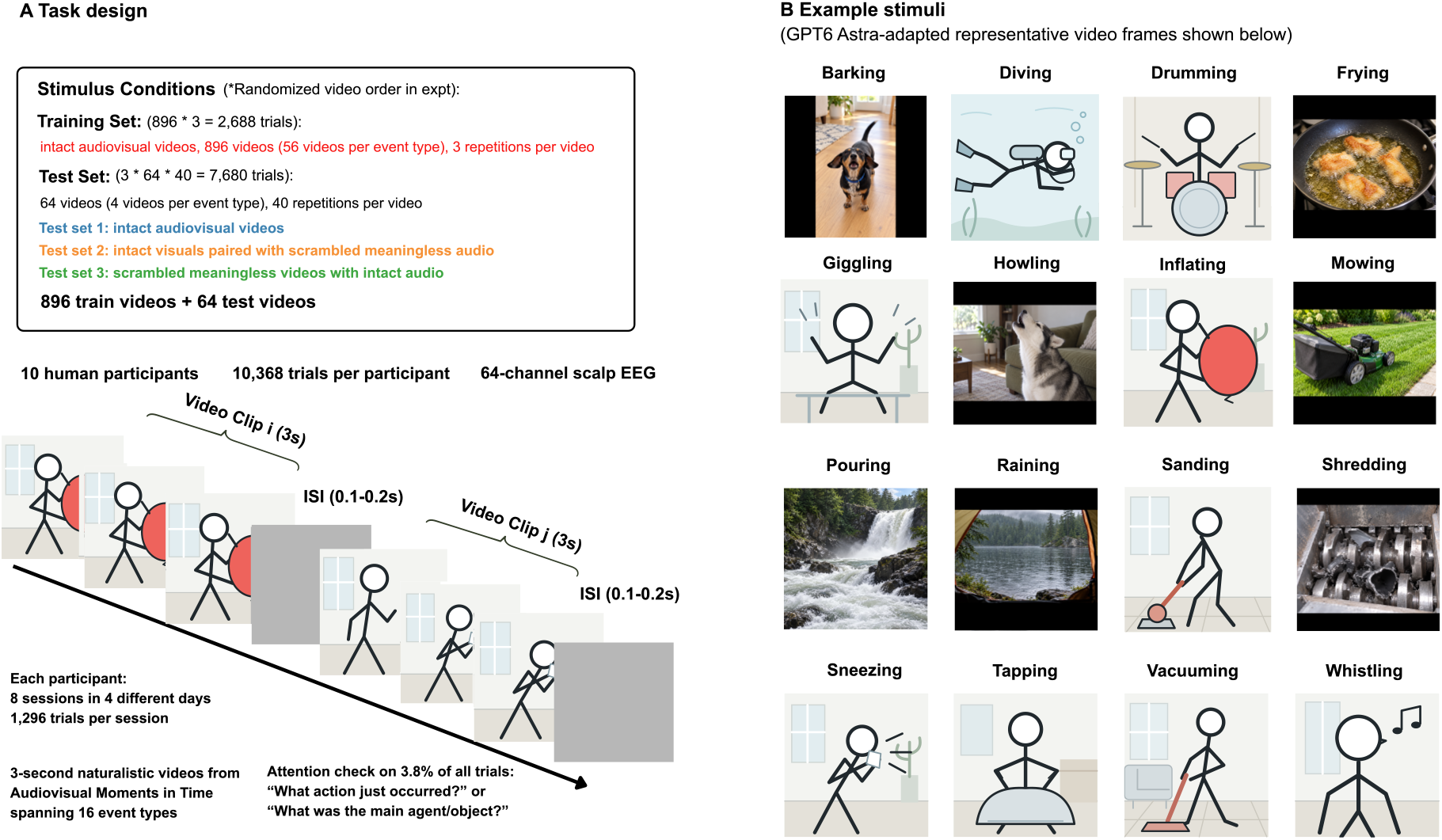
Experimental task design and example audiovisual stimuli. (A) Task design of the experiment. Participants freely viewed 3-s naturalistic audiovisual clips, each followed by a randomly jittered inter-stimulus interval (ISI) of 0.1–0.2 s. For illustration purposes, only part of the trial sequence is shown. (B) Illustrative examples from each of the 16 event categories. To address licensing concerns, the original video frames in both panels have been replaced with GPT6 Astra-adapted illustrations preserving the depicted event content. These illustrations were not presented to participants.

In the present study, we used the curated AVMIT stimulus set comprising 960 three-second videos from 16 event categories, with 60 videos per category. We used these 960 three-second videos as the intact audiovisual stimulus pool in the present dataset. The 16 event categories were barking, diving, drumming, frying, giggling, howling, inflating, mowing, pouring, raining, sanding, shredding, sneezing, tapping, vacuuming, and whistling. For each category, 56 videos were assigned to the training set and the remaining four videos to the test set, yielding 896 training videos and 64 test videos in total. Training and test videos were therefore distinct exemplars of the same 16 event categories.

The same 64 held-out video exemplars were used to construct three test sets that differed in the availability of coherent visual and auditory event information:

In Test set 1, the original audiovisual videos were presented without modification.

In Test set 2, the visual stream was preserved while the accompanying audio was temporally scrambled. Specifically, the original audio track was extracted at a sampling rate of 44.1 kHz, and the temporal order of the audio samples was randomly permuted. The scrambled audio was then recombined with the unchanged visual stream, preserving the duration and constituent audio samples while disrupting the meaningful temporal structure of the soundtrack.

In Test set 3, the original audio track was preserved while the visual stream was scrambled. Simply randomizing the temporal order of video frames would not necessarily eliminate meaningful event information, because individual frames could still contain recognizable objects, agents, scenes, or action-related cues. We therefore additionally disrupted the spatial structure of each frame while retaining basic low-level visual information. Specifically, each frame was independently phase-scrambled in the Fourier domain, preserving its Fourier amplitude spectrum and thus much of its low-level spatial-frequency content while randomizing phase information and disrupting recognizable image structure. The resulting phase-scrambled frames were also presented in a randomized temporal order, further disrupting coherent motion and event dynamics. This scrambled visual stream was then recombined with the original intact audio track.

Thus, the three test sets contained the same underlying video exemplars but differed systematically in the availability of coherent modality-specific event information: intact audiovisual information (Test set 1), intact visual information with scrambled audio (Test set 2), and intact auditory information with scrambled visual input (Test set 3). This design enabled the same event categories and video identities to be evaluated under different audiovisual conditions while minimizing meaningful information in the scrambled modality.

### Experiment

In the present experiment, we used 16 action categories, each containing 60 audiovisual clips of 3 s duration, resulting in 960 unique clips. The 60 clips in each category were divided into two subsets, with 56 clips assigned to the training set and 4 clips assigned to the test set. Thus, the training set contained 896 clips and the test set contained 64 clips.

To dissociate the contributions of visual and auditory information to semantic processing, each clip in the test set was presented under three conditions. In the intact audiovisual condition, the original audiovisual clip was presented, with both the visual and auditory streams conveying meaning. intact visual condition with scrambled audio, the original visual stream was preserved, while the original sound was replaced by noise generated by randomly shuffling the audio samples, rendering the auditory input meaningless while preserving meaningful visual input. In the intact auditory condition with scrambled visual input, the original audio stream was preserved, whereas the visual stream was disrupted by Fourier phase scrambling and temporal frame shuffling, removing recognizable visual content and coherent motion while retaining basic low-level visual properties.

The experiment used a mixed design. Training-set trials and trials from the three test-set conditions were randomly interleaved within the same session, and all conditions were balanced within each session. Each participant completed 8 formal sessions. In the training set, each clip was repeated 3 times, yielding 2,688 training trials. In the test set, each of the 64 clips was presented 40 times in each of the three conditions, yielding 7,680 test trials. The full experiment contained 10,368 trials. Each trial presented a 3-s video stimulus, followed by a random inter-stimulus interval (ISI) of 0.1-0.2s. The total stimulus presentation time was approximately 9.2 h, completed across eight sessions over four different days.

The experiment was implemented in Python 3.10 using PsychoPy version 2024.2.4^14^. Stimuli were presented on a DELL monitor with a resolution of 1920 × 1080 pixels and a screen width of 54 cm. Participants viewed the screen from a distance of approximately 60 cm. All clips were presented centrally on a gray background in a square format, with a side length of approximately 8.4° of visual angle. To maintain attention and engagement, 5% of trials in the training set and in test-set conditions 1 and 2 included a detection task related to the video content. Participants freely viewed the clips throughout the experiment. Natural eye movements were allowed, no fixation was required, and participants could pause and rest between trials by pressing the space bar.

### EEG data recording

Continuous EEG signals were recorded while participants viewed the clips using an ANT Neuro eego− mylab system. Conductive gel was used to keep electrode impedances below 10 kΩ whenever possible. EEG signals were acquired with a 64-channel EEG system, with electrode positions arranged according to the extended international 10-10 system. The online reference electrode was Cz. Signals were digitized at a sampling rate of 1000 Hz. At the onset of each clip, a 10-ms event trigger pulse was sent to the EEG acquisition system through a parallel port. During EEG recording, ECG signals were simultaneously collected from each participant. The same 10-ms event trigger pulse was sent to the ECG acquisition system, allowing ECG data to be aligned with EEG on a trial-by-trial basis.

### EEG data preprocessing

For basic quality control and technical validation, EEG data were preprocessed using an MNE-Python^15^ based pipeline. Raw BrainVision EEG data were imported, EOG channels and a custom electrode montage were specified. Continuous data were band-pass filtered from 0.03 to 50 Hz, notch-filtered at 48–52 Hz, and downsampled to 100 Hz. Bad channels identified using PyPREP^16^ were interpolated. Epochs were extracted from −1.5 to 5 s relative to video-stimulus onset and baseline-corrected using −0.1 to 0 s.

Data from eight sessions were concatenated. A 0–3 s copy was used for quality control and independent component analysis (ICA)^17^ fitting after 1-Hz high-pass filtering. ICA components were identified using EOG/ECG information and manual inspection, then removed from the full −1.5 to 5 s epochs. Data were re-referenced to the average of two mastoid electrodes (M1 and M2). Bad epochs for the primary analysis were rejected based on the 0–3 s interval. Preprocessed data and processing logs were saved in FIF and NumPy formats for subsequent time-resolved decoding analyses.

## Data Records

### Stimuli

The released stimuli are stored in the Videos_for_exp directory and comprise 1,088 three-second MP4 videos. The training set (Exp1) contains 896 intact audiovisual videos, with 56 exemplars from each of 16 event categories. The test set (Exp2) comprises three conditions (Cond1–Cond3), each containing the same 64 held-out exemplars, for a total of 192 videos.

Training files are organized under Videos_for_exp/Exp1/type1–type16, with filenames ranging from video_1.mp4 to video_56.mp4. Test files are organized under Videos_for_exp/Exp2/Cond1–Cond3 and the corresponding category directories, with filenames ranging from video_1.mp4 to video_4.mp4. The category-label mapping is provided in Exp1/Type-naming-list.txt. Corresponding files across the three test conditions share the same category and exemplar identifiers.

### BIDS-formatted data

The BIDS_Data directory contains the continuous EEG, ECG, and behavioral data organized according to EEG-BIDS version 1.9.0 using MNE-BIDS version 0.18.0^18^. The BIDS-formatted dataset was checked using the BIDS Validator, which reported no validation errors. The dataset includes 10 participants, each with eight recording sessions (ses-01– ses-08), yielding 80 participant-session recordings. Within each session, continuous EEG and concurrently acquired ECG recordings, together with their associated metadata, are stored in the eeg directory. EEG data are provided in BrainVision format, whereas ECG data are provided as BIDS-compatible physiological recordings named_recording-cardiac_physio.tsv.gz. Trial-level behavioral records are stored in the beh directory. The corresponding _scans.tsv file indexes the recording files included in each session. The organization of the BIDS-formatted dataset is illustrated in Fig. 2.

**Fig. 2.**
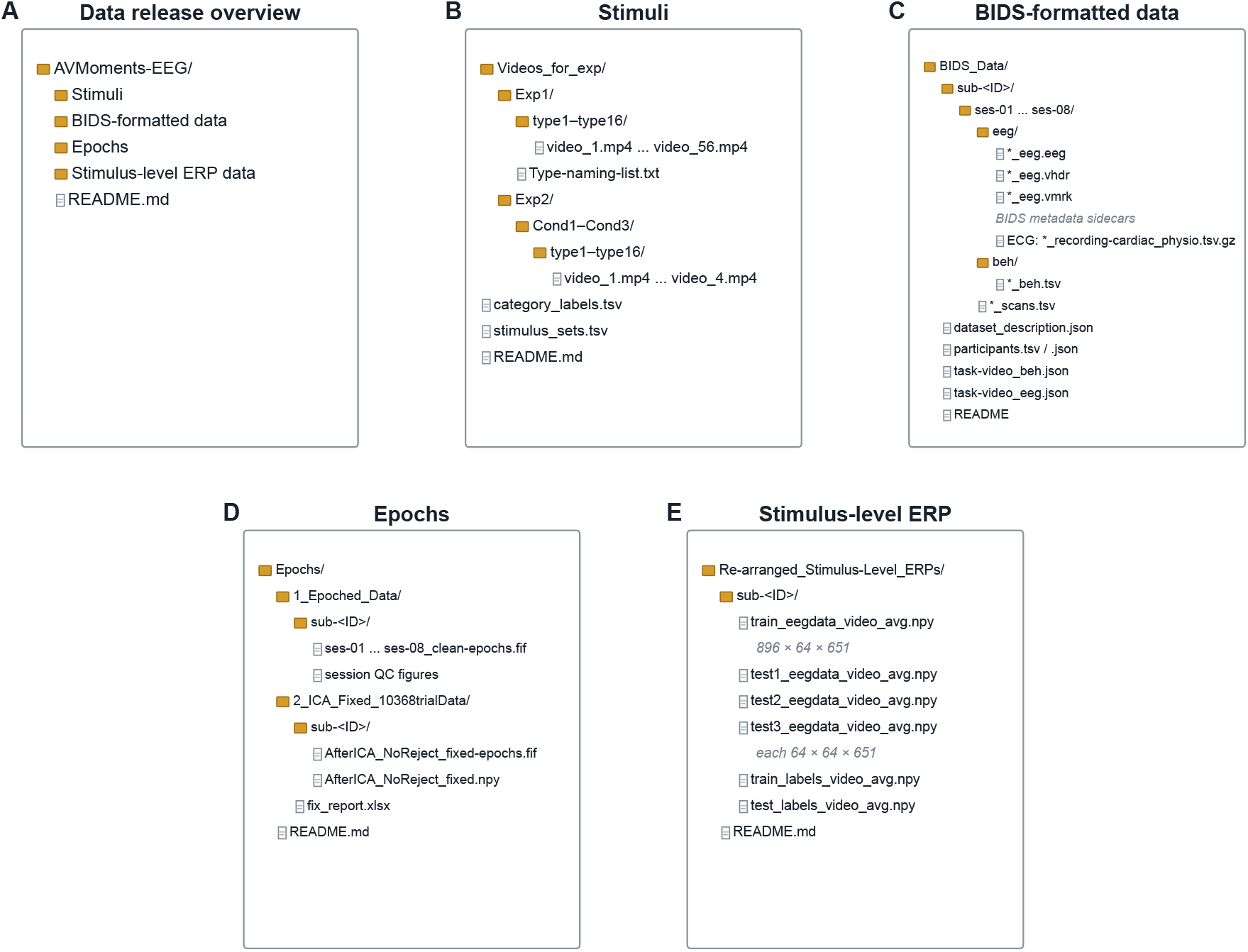
Organization of the AVMoments-EEG data records. (A) Overview of the four data components. (B) Stimulus videos organized into training and test sets. (C) BIDS-formatted data organized by participant, session, and modality. (D) Session-level and participant-level epoch derivatives. (E) Stimulus-level ERP arrays reorganized and averaged by video identity.

The EEG sidecar files provide recording parameters, channel descriptions, electrode coordinates, and coordinate-system information. Event timing and labels are provided in_events.tsv and defined in _events.json. Because the marker naming convention for the first two participants differed from that used for the remaining participants, event labels were harmonized during BIDS conversion. The _events.tsv files should therefore be used as the primary reference for event timing and codes. Dataset-level metadata and participant demographics are provided in dataset_description.json, participants.tsv, participants.json, and README. These BIDS-formatted continuous recordings constitute the source data from which the preprocessed epochs and stimulus-level ERP arrays were generated.

### Epochs

Epoch-level derivatives for the final cohort of 10 participants are provided at two preprocessing stages. The 1_Epoched_Data directory contains session-level epochs before ICA-based artifact correction. Each participant folder contains eight FIF files, named ses-01_clean-epochs.fif through ses-08_clean-epochs.fif, together with quality-control figures showing the continuous recordings and session-averaged evoked responses. Preprocessing and epoch construction are described in Methods. Each epoch spans −1.5 to 5.0 s relative to video onset and contains 64 channels and 651 time points at 100 Hz. The experimental design specified 1,296 trials per session and 10,368 trials across eight sessions. The number of available session-level epochs occasionally differed from the expected count because of missing or duplicated acquisition triggers or because some events occurred too close to a recording boundary to yield a complete epoch.

The 2_ICA_Fixed_10368trialData directory contains participant-level post-ICA epochs before trial-level artifact rejection. For each participant, epochs from the eight sessions were concatenated and processed using the ICA correction and re-referencing procedures described in Methods. Each participant is represented by one MNE FIF file and one NumPy array, both identified by AfterICA_NoReject_fixed in the filename. The NumPy arrays have dimensions of 10,368 trials × 64 channels × 651 time points, corresponding to trial × channel × time, and are stored as 64-bit floating-point values. The NoReject designation indicates that ICA correction has been applied but that the trial-level peak-to-peak rejection used to construct the stimulus-level ERP data has not yet been performed.

To provide a common trial axis across participants, the post-ICA epochs were aligned to a fixed grid of 10,368 trial positions. Nineteen positions for which no acquired EEG epoch was available are represented by all-zero placeholder epochs labelled with the event code s9999: 17 for sub-008ckp, 1 for sub-010lxc, and 1 for sub-013zzy. These placeholders do not represent recorded EEG signals and must be excluded before subsequent analyses. Their locations and all trial-grid corrections are documented in fix_report.xlsx.

### Re-arranged stimulus-level ERP data

To facilitate stimulus-level analyses, we additionally provide EEG responses reorganized and averaged according to individual video identity. These files were generated from the preprocessed epoch-level EEG data after ICA correction. As an additional quality-control step, we calculated the peak-to-peak amplitude for each channel within the 0–3 s video-presentation period of every epoch. Channels with a peak-to-peak amplitude exceeding 300 μV were flagged, and epochs in which at least 32 of the 64 recorded channels exceeded this threshold were excluded. The remaining epochs were matched to the corresponding behavioral records using the stimulus condition, event type, and video number.

For each participant, retained epochs were first separated into the training set and the three test conditions. The training set comprised 896 unique intact audiovisual videos, consisting of 56 videos from each of the 16 event categories. Each test set comprised the same 64 test video identities, with four videos from each event category, presented under one of three audiovisual conditions: intact audiovisual videos (Test set 1), intact visual videos paired with scrambled audio (Test set 2), or scrambled visual videos paired with intact audio (Test set 3). For each unique video within each condition, all retained repetitions were averaged to obtain a single stimulus-level ERP response.

The resulting files are stored separately for each participant as train_eegdata_video_avg.npy, test1_eegdata_video_avg.npy, test2_eegdata_video_avg.npy, and test3_eegdata_video_avg.npy. The training array has dimensions 896 × 64 × 651, whereas each test array has dimensions 64 × 64 × 651, corresponding to stimulus × channel × time, respectively. The 651 time points span −1.5 to 5 s relative to video onset at a sampling rate of 100 Hz. Stimuli are ordered first by event category and then by video identity within each category. Corresponding event-category labels are provided in train_labels_video_avg.npy and test_labels_video_avg.npy, with dimensions 896 and 64, respectively. Training labels contain 56 consecutive entries for each of the 16 event categories, whereas test labels contain four consecutive entries for each category.

## Technical Validation

### Behavioral performance

To verify that participants remained attentive throughout the extended EEG recording sessions, attention-check questions were presented on approximately 3.8% of trials (390 of 10368 trials). Responses were scored using a keyword-matching procedure against the action category and the main agent or object in each video. Participants were asked either to identify the action that had just occurred or to report the main agent or object in the preceding video. Performance was consistently high across participants, with a mean accuracy of 81.1 ± 8.3%, indicating that participants remained engaged with the video content throughout the experiment.

### Stimulus-evoked EEG responses

We first examined whether the processed stimulus-level EEG data retained robust stimulus-evoked temporal and spatial structure. For each participant, responses were averaged across stimuli within each of the four stimulus sets and were subsequently averaged across participants. Analyses and visualizations were based on the 61 scalp EEG electrodes, excluding the EOG channel and the two mastoid electrodes (M1 and M2).

Grand-average ERPs showed clear stimulus-locked responses across the scalp for the training set and all three test sets (Fig. 3A). Following video onset, prominent voltage deflections emerged within the first several hundred milliseconds and were followed by sustained activity throughout the 3-s viewing period. The overall temporal profiles were broadly similar across the training and test sets, demonstrating that robust evoked responses were preserved after preprocessing and stimulus-level averaging. The spatially colored butterfly plots further illustrate the contribution of electrodes distributed across the scalp to these response profiles.

**Figure 3.**
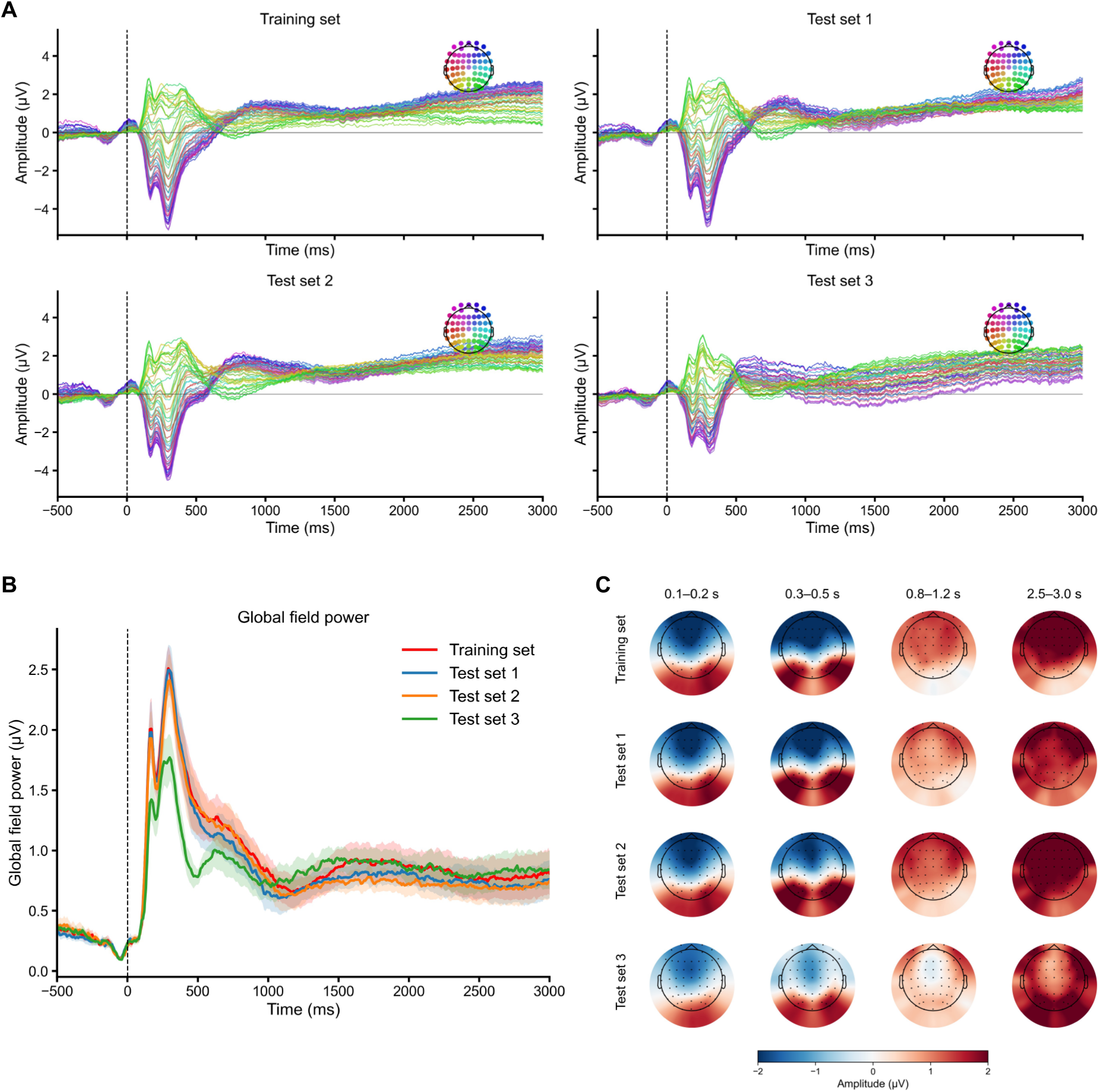
Stimulus-evoked temporal and spatial characteristics of the EEG responses. (A) Grand-average ERP butterfly plots for the training set and the three test sets. For each participant, stimulus-level responses were first averaged across videos within each set and subsequently averaged across the 10 participants. Each colored trace represents one of the 61 scalp EEG electrodes; colors correspond to electrode locations shown in the scalp inset. The EOG channel and the two mastoid electrodes (M1 and M2) were excluded. The vertical dashed line indicates video onset (0 ms). (B) GFP for the four stimulus sets, calculated for each participant as the spatial standard deviation across the 61 scalp electrodes at each time point. Lines indicate the group mean and shaded regions indicate ±SEM across participants. (C) Group-average scalp voltage distributions during four representative time windows: 100–200 ms, 300–500 ms, 800–1200 ms, and 2500–3000 ms.

We next quantified the overall strength of the scalp electric field using global field power (GFP), calculated separately for each participant as the spatial standard deviation of voltage across the 61 scalp electrodes at each time point. GFP increased sharply following stimulus onset and reached its largest values within the first several hundred milliseconds, followed by sustained field strength throughout the remainder of the video presentation (Fig. 3B). Although the magnitude and temporal profile varied somewhat across stimulus sets, all four sets exhibited clear stimulus-evoked increases in GFP relative to the prestimulus period.

Finally, we characterized the spatial distribution of the evoked responses over four representative temporal intervals spanning early and sustained stages of video processing: 100–200 ms, 300–500 ms, 800–1,200 ms, and 2,500–3,000 ms (Fig. 3C). The scalp maps revealed structured and temporally evolving voltage distributions across all four stimulus sets. Together, the ERP waveforms, GFP time courses, and scalp topographies demonstrate that the released stimulus-level EEG responses preserve robust temporal and spatial structure throughout naturalistic video viewing.

### Classification-based EEG decoding of event-category information

We next assessed whether the processed EEG responses retained information that could support multivariate analyses of naturalistic event representations. The 16 event categories were evaluated using pairwise classification, yielding 120 event-pair decoding problems. Classifiers were trained on EEG responses to the 896 intact audiovisual videos in the training set and tested on the 64 independent videos in each of the three test sets. Importantly, the training and test sets contained distinct video exemplars. Thus, above-chance performance reflects generalization of event-related neural information to previously unseen videos rather than discrimination of repeated presentations of identical stimuli.

Time-resolved decoding revealed robust event-category information for both the intact audiovisual test set (Test set 1) and the intact-visual/scrambled-audio test set (Test set 2) (Fig. 4A). Decoding accuracy remained close to the 50% chance level before stimulus onset, increased rapidly after video onset, reached approximately 60% at its early peak, and remained above chance over extended portions of the video presentation. In contrast, decoding for the scrambled-visual/intact-audio condition (Test set 3) was substantially weaker and remained close to chance over most of the time course.

**Figure 4.**
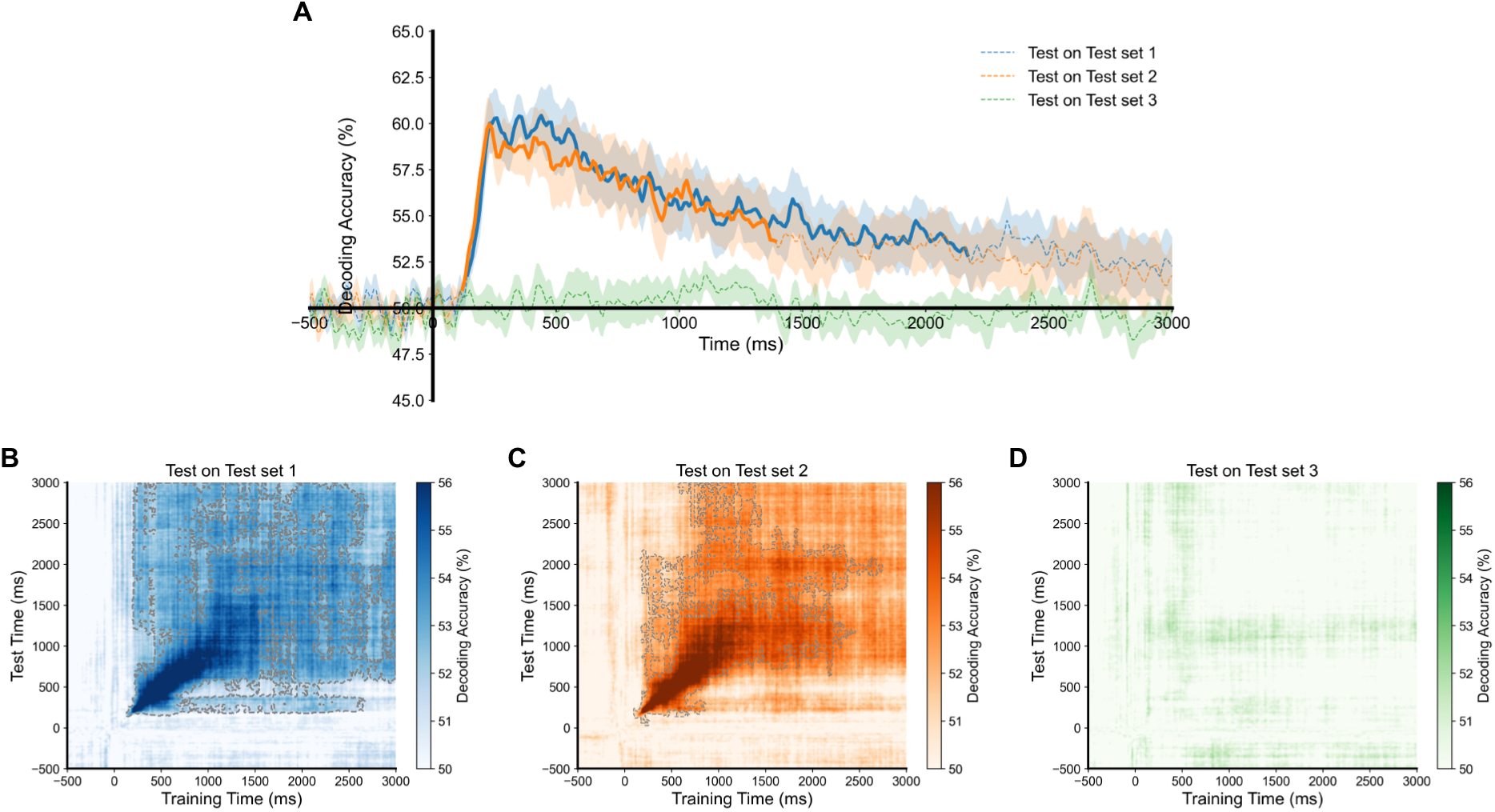
EEG decoding and temporal generalization of event-category information. (A) Time-resolved and cross-temporal pairwise decoding of the 16 event categories. Classifiers were trained on stimulus-level EEG responses to the 896 intact audiovisual videos in the training set and independently tested on the 64 videos in each test set: intact audiovisual videos (Test set 1), intact visual videos paired with scrambled audio (Test set 2), and scrambled visual videos paired with intact audio (Test set 3). Decoding accuracy was averaged across the 120 pairwise event comparisons and across participants. The horizontal line indicates the 50% chance level. Shaded regions indicate ±1 SEM across participants. Statistical significance against chance was assessed using cluster-based permutation tests^20^; solid portions of each curve indicate significant temporal clusters, whereas dashed portions indicate non-significant time points. The vertical solid line indicates video onset. (B-D) Cross-temporal event-category decoding for the same three test sets: (B) test on test set 1, (C) test on test set 2, and (D) test on test set 3. Classifiers trained at each training-set time point were evaluated across all test-set time points. Statistical significance against the chance level was assessed using cluster-based permutation tests across the two-dimensional training-time × test-time space. Gray dashed contours delineate significant clusters.

Cross-temporal decoding further characterized the temporal organization and persistence of event-discriminative EEG patterns^19^. For Test sets 1 and 2 (Fig. 4B-C), decoding generalized strongly along the training-time/test-time diagonal and also extended to off-diagonal time points, indicating that event-discriminative patterns were not confined to isolated moments but showed temporal persistence and generalization. This structure was markedly weaker for Test set 3 (Fig. 4D). Together, the time-resolved and cross-temporal analyses demonstrate that the dataset contains temporally resolved and generalizable information about naturalistic event categories. In particular, event-discriminative EEG patterns learned from an independent audiovisual training set generalized robustly to unseen test videos when their visual content remained intact, providing an information-based validation of the utility of the released dataset for multivariate decoding and representational analyses. All classification-based EEG decoding were implemented based on NeuroRA^20^.

## Data and Materials Availability

The AVMoments-EEG dataset is publicly available on Zenodo at https://doi.org/10.5281/zenodo.22227356. Due to licensing restrictions, access to the video stimuli used in this study can be obtained by contacting the corresponding authors.

## Code Availability

the code used to run the experiment, preprocess the EEG recordings, and validate the released data is publicly available on GitHub at https://github.com/ShuningTangg/AVMoments-EEG.

## Author Contributions

Zitong Lu and Dongwei Li conceptualized the study. Shuning Tang programmed the experiment. Leiting Li collected the data. Shuning Tang and Zitong Lu processed the data, performed statistical analyses, validated the results, curated and shared the data, and prepared the figures. Zitong Lu and Dongwei Li supervised the study. All authors contributed to manuscript writing.

